# The primary motor cortex prospectively computes the spinal reflex

**DOI:** 10.1101/2020.06.29.170928

**Authors:** Tatsuya Umeda, Tadashi Isa, Yukio Nishimura

**Author notes:** Correspondence to (T.U.), (Y.N.).

## Abstract

The spinal reflex transforms sensory signals to generate muscle activity. However, it is unknown how the motor cortex (MCx) takes the spinal reflex into account when performing voluntary limb movements. We simultaneously recorded the activity of the MCx, afferent neurons, and forelimb muscles in behaving monkeys. We decomposed muscle activity into subcomponents explained by the MCx or afferent activity using linear models. Long preceding activity in the MCx, which is responsible for subsequent afferent activity, had the same spatiotemporal contribution to muscle activity as afferent activity, indicating that the MCx drives muscle activity not only by direct descending activation but also by trans-afferent descending activation. Therefore, the MCx implements internal models that prospectively estimate muscle activation via the spinal reflex for precise movement control.

## Main Text

How the central nervous system generates appropriate muscle activity for achieving intended movements is a long-standing question in the field of movement control (*1, 2*). Extensive investigation has shown that various movement-related parameters are represented by neural activity in the primary motor cortex (M1) (*3-9*), but scientists still have not reached a consensus on what information the M1 sends to spinal motoneurons. A new perspective has been proposed recently in which the neural population in the M1 acts like a dynamical system (*10*). However, this concept has not answered the question of how the population dynamical state in the M1 has evolved to drive muscle activity (*11*). Despite the fact that somatosensory inputs from peripheral afferents contribute substantially to the generation of muscle activity during voluntary movements (*12-15*), these prior studies have failed to evaluate the effect of the spinal reflex on information flow from the M1 to spinal motoneurons. Thus, a conceptual framework of this information flow should be derived from an investigation of the movement control systems that incorporate supraspinal structures, such as the M1, and the spinal reflex arc, which is composed of muscles and peripheral afferents.

In this study, we simultaneously recorded electrocorticographic signals from the motor cortex (MCx), including the M1 and the dorsal (PMd) and ventral (PMv) premotor cortex, the activity of a population of lower cervical afferents, and electromyographic signals from the forelimb muscles in two monkeys when performing reaching and grasping movements (Fig. 1, A–C). We first examined how descending inputs from the MCx (descending input) and inputs from peripheral afferents (afferent input) are involved in generating muscle activity (Fig. 1D). Neural ensemble activity in the M1 has been shown to represent muscle activity using a linear model (*16, 17*). We built a linear model using descending and afferent inputs to account for the subsequent muscle activity. The model reconstructed the overall temporal pattern of muscle activity more accurately than a model built from surrogate shuffled data (fig. S1) and models built from each input alone (Fig. 1, E and F). These results suggest that afferent input is necessary information for the reconstruction of muscle activity. We assessed how each descending and afferent input contributed to the reconstructed muscle activity by calculating each subcomponent of the reconstructed activity (Fig. 2A). While the descending component increased before movement onset, similar to the observed muscle activity, the afferent component increased only after movement onset (Fig. 2, B and C). The sequential contribution of descending and afferent inputs on muscle activity was also derived from analyses by building linear models in sliding time windows (fig. S2). Thus, premovement muscle activity could be explained by descending input, while that after movement onset could be explained by sum of descending and afferent inputs.

**Fig. 1.**
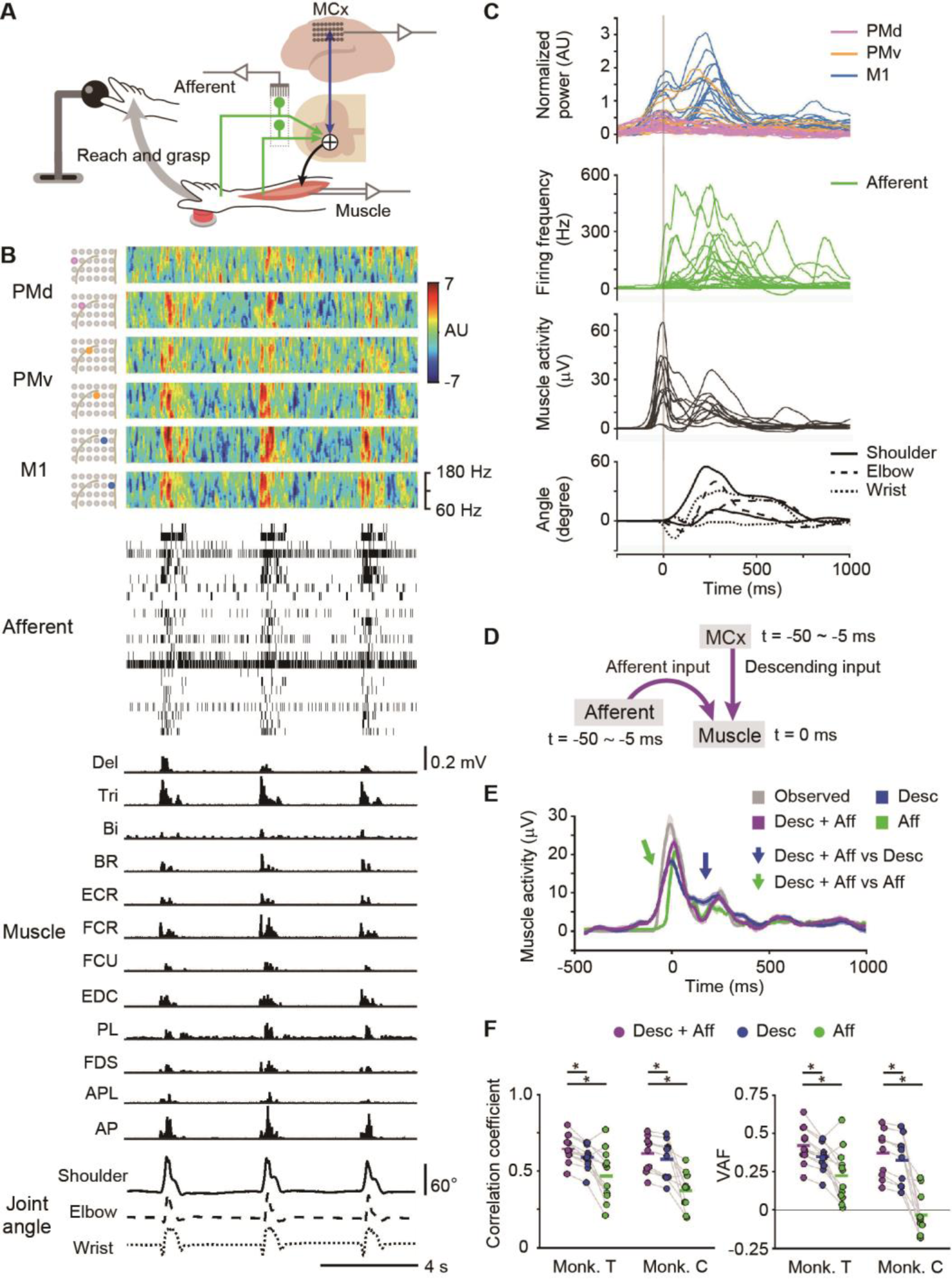
Simultaneous recording of cortical and peripheral activity. (**A**) Schematic of the experimental setup. (**B**) An example of simultaneous recording of three trials by monkey T. *Top*: Power spectrograms in the MCx. *Second*: Raster plots of peripheral afferent activity. *Third*: Activity of forelimb muscles. *Bottom*: Forelimb joint angles along the extension-flexion axis. (**C**) Modulations of cortical and peripheral activity in monkey T aligned to movement onset. *Top*: High-gamma activity. *Second*: Instantaneous firing rate of peripheral afferents. *Third*: Forelimb muscles. *Bottom*: Joint angles. Vertical line, movement onset. (**D**) Model to account for muscle activity from descending and afferent inputs. (**E**) Average modulation of the observed muscle activity, reconstruction using descending and afferent inputs, and each input aligned to movement onset. Shaded areas, SEM. Arrows, different points between two models. (**F**) Correlation coefficient and variance accounted for (VAF) between the observed and reconstructed traces (monkey T, n = 12 muscles; monkey C, n = 10 muscles; *P* < 0.005, one-way analysis of variance [ANOVA], \**P* < 0.0005, paired t-test). Superimposed bars, mean.

**Fig. 2.**
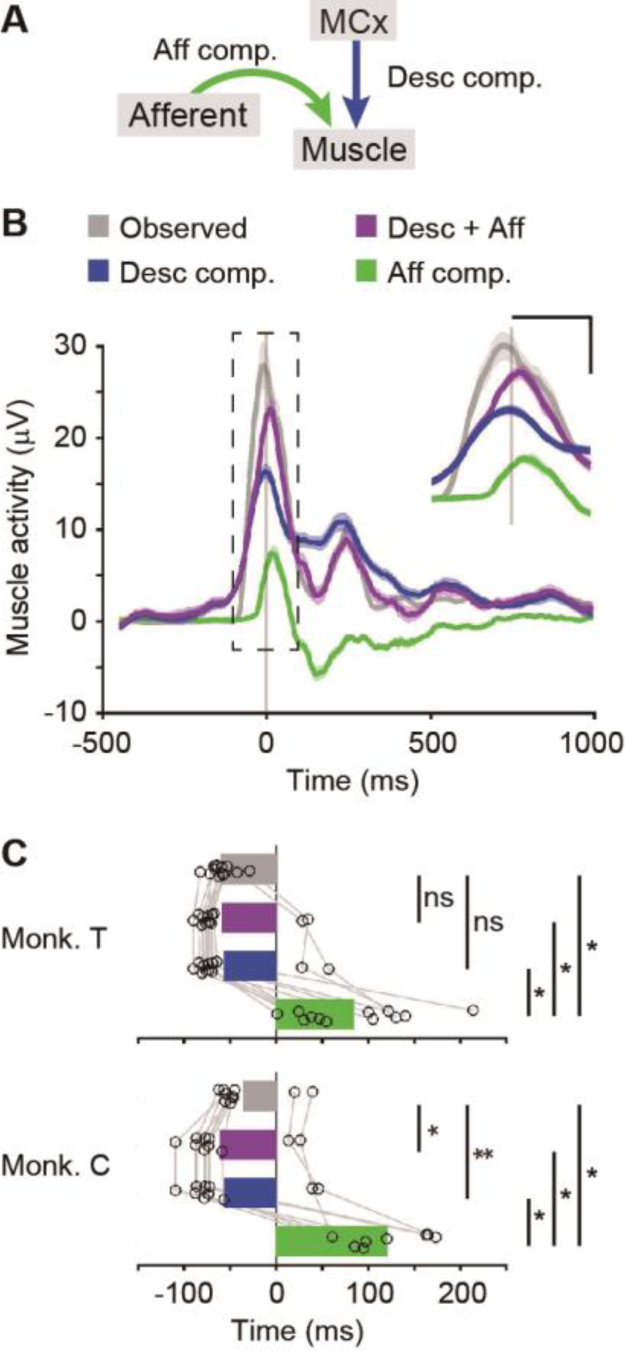
Sequential integration of descending and afferent inputs. (**A**) Model to account for muscle activity from descending and afferent inputs. (**B**) Average modulation of the observed muscle activity, reconstruction using descending and afferent inputs, and each subcomponent aligned to movement onset. Vertical lines, movement onset. Shaded areas, SEM. Scale bars, 100 ms and 10 μV. (**C**) Onset times of the observed muscle activity, of the reconstruction using descending and afferent inputs, and of each subcomponent (monkey T, n = 12 muscles; monkey C, n = 10 muscles; *P* < 10^−9^, one-way ANOVA, \**P* < 0.0005, \*\**P* < 0.05, paired t-test). ns, not significant. Superimposed bars, mean.

We next examined how descending or afferent inputs contributed to the reconstruction of the activity of each muscle. The descending component during a period in which the movement-related modulation of muscles was observed had positive values for all muscles (Fig. 3, A and B), indicating that the descending input has a facilitatory effect on all muscles. Conversely, the afferent input exhibited facilitatory and suppressive effects (Fig. 3, A and B). Since limb movement modulates activity in most peripheral afferents, we investigated whether the wide variety of afferent components might be explained by the response to limb movement. When the monkeys began to reach, monkey T flexed the wrist joint only (fig. S3A). Immediately after the initial movement, the afferent input exerted a facilitatory effect on a wrist extensor muscle (extensor carpi radialis) and a suppressive effect on wrist flexor muscles (flexor carpi radialis [FCR], flexor digitorum superficialis) (fig. S3, B and C). These effects on extensor and flexor muscles could be explained by the stretch reflex and reciprocal inhibition, respectively. The results from monkey C also show a similar relationship between the afferent component and initial joint kinematics (fig. S3). Thus, the effects of afferent inputs on muscle activity are at least partly accounted for by the action of the spinal reflex.

**Fig. 3.**
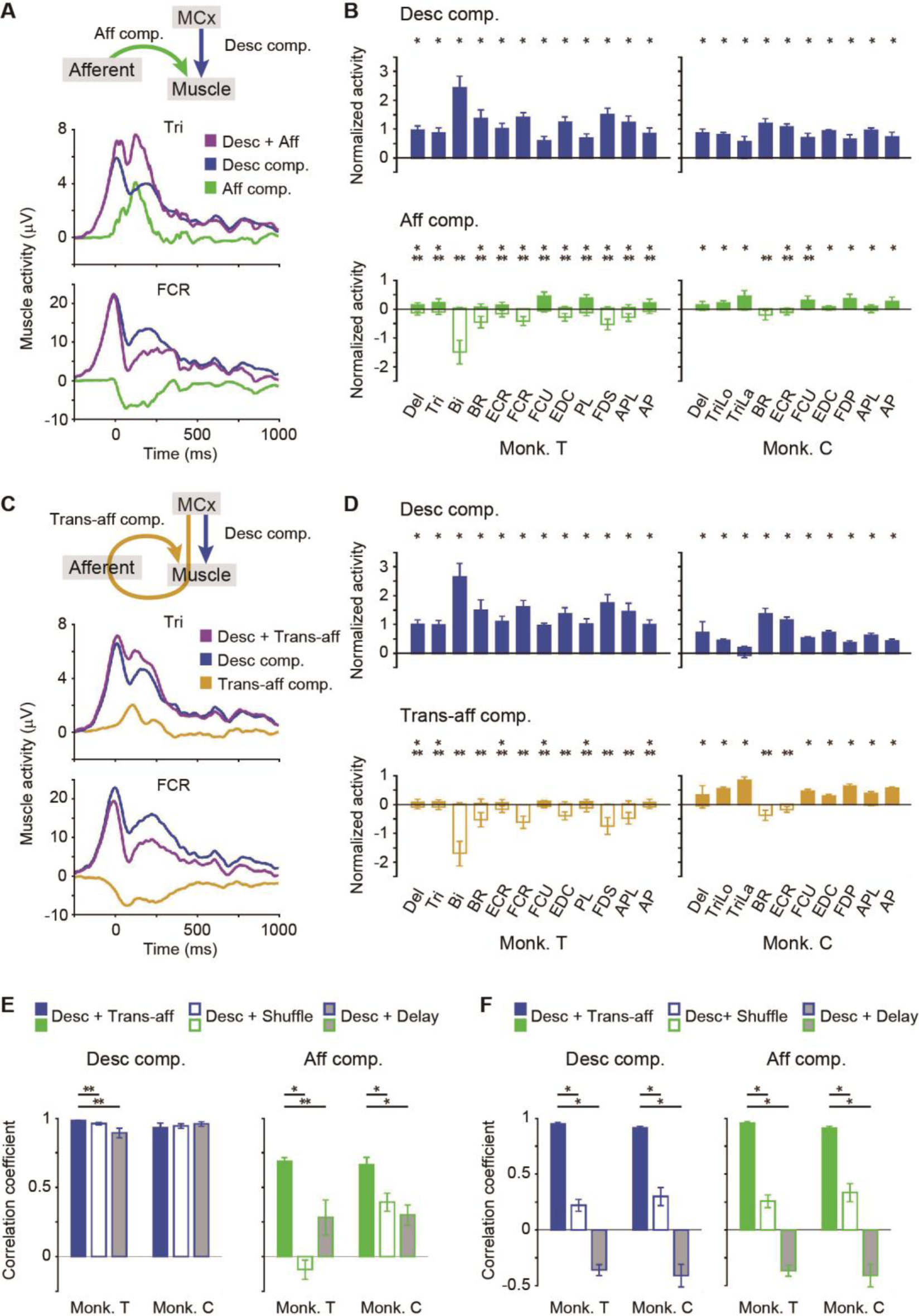
MCx encodes the activation of muscles via the spinal reflex. (**A**) Average modulation of the reconstruction using descending and afferent inputs, and each subcomponent aligned to movement onset. (**B**) Modulation of cortical and afferent components for each muscle (monkey T, n = 21 sessions; monkey C, n = 7 sessions; \**P* < 0.05, unpaired t-test for positive value, \*\**P* < 0.05, unpaired t-test for negative value). Error bars, SD. (**C, D**) Same as (A, B) for descending and trans-afferent inputs. (**E**) Temporal similarity of the descending or afferent component between models (monkey T, n = 12 muscles; monkey C, n = 10 muscles; *P* < 0.05 for both components of monkey T and afferent component of monkey C, one-way ANOVA, \**P* < 0.001, \*\**P* < 0.05, paired t-test). Superimposed bar graphs and error bars, mean ± SEM. (**F**) Same as (E) for the spatial similarity of the descending or afferent component across muscles between models (*P* < 0.05, one-way ANOVA, \**P* < 0.001, paired t-test).

Taken together, our results suggest that the descending input from the MCx drives the premovement activity of muscles, which subsequently affects muscle activity through the spinal reflex arc. Since the activity in the MCx also accounts for the subsequent afferent activity 60-70 ms later (fig. S4), we hypothesized that the activity in the MCx drives muscle activity through the descending and spinal reflex pathways (trans-afferent input) in addition to the descending pathway (descending input) (Fig. 3C). To test this hypothesis, we built a linear model using descending and trans-afferent inputs to account for the subsequent muscle activity. The model reconstructed the overall temporal pattern of muscle activity more accurately than the model built from the descending input alone (fig. S5). The results suggest that trans-afferent input is necessary for the precise reconstruction of muscle activity.

We next examined how the trans-afferent input contributed to the reconstructed muscle activity by calculating its subcomponent. The temporal profile of the trans-afferent component shown in Fig. 3C was similar to that of the afferent component of a model obtained from the descending and afferent inputs shown in Fig. 3A. Furthermore, the distribution of the size of the afferent component across muscles was similar to that of the trans-afferent component (Fig. 3, B and D). To evaluate these similarities, we used surrogate shuffled data of the trans-afferent input or delayed activity in the MCx as a control for trans-afferent input and calculated the corresponding subcomponent (shuffled or delayed component, respectively) (fig. S6). The similarity of the temporal profile (temporal similarity) and the similarity of the distribution across muscles (spatial similarity) between the afferent and trans-afferent components was higher than the temporal and spatial similarity between the afferent and shuffled components and between the afferent and delayed components, respectively (Fig. 3. E and F). Thus, long-preceding activity in the MCx had the same spatiotemporal contribution to muscle activity as afferent activity. These results suggest that the MCx affects muscle activity by trans-afferent activation through the descending and spinal reflex pathways in addition to direct activation via the descending pathway.

Finally, we asked to what extent the activity in individual areas of the MCx contributed to the reconstruction of muscle activity. We examined weight values assigned to four types of cortical input (Fig. 4A). For all four inputs, the weight values given to M1 activity were larger than those given to PMd or PMv activity (Fig. 4B). The M1 electrodes with the highest weight value for trans-afferent input were located just anterior to the central sulcus, as for the descending input (Fig. 4A). Thus, muscle activity could mostly be accounted for by the trans-afferent input from a subset of M1 regions, rather than the PMd and PMv. These results suggest that the M1 implements an internal model that prospectively computes muscle activation via the spinal reflex to solve an inverse problem for executing intended movements.

**Fig. 4.**
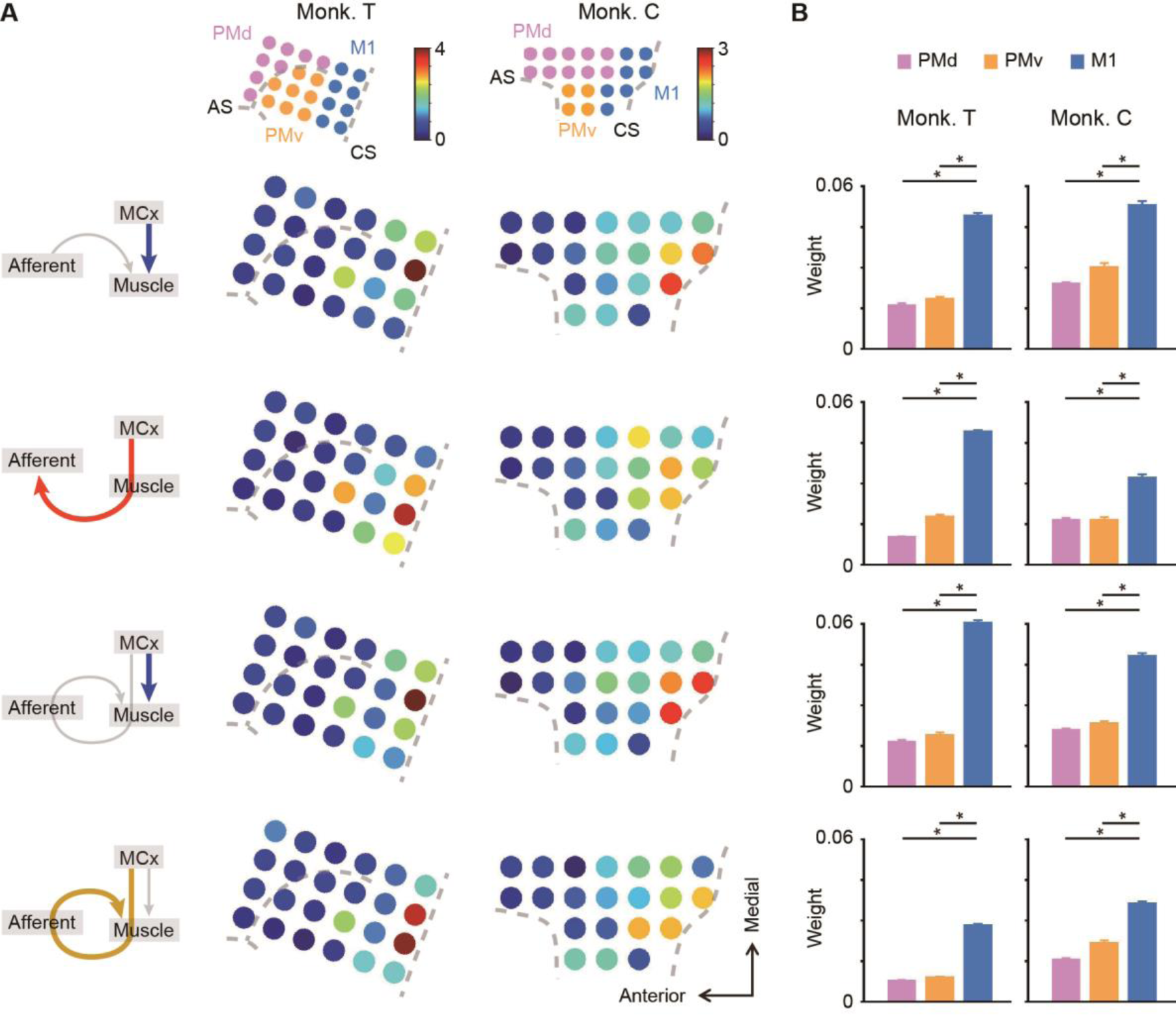
The M1 is a major predictor of muscle activity. (**A**) Color maps representing the average weight values of MCx activity at each electrode in models that predict muscle activity. Thick colored arrows in the left schematics show the inputs for which the weight values are represented. CS, central sulcus; AS, arcuate sulcus. (**B**) The average weight values of each cortical area for the prediction of muscle activity (monkey T, n = 12 muscles; monkey C, n = 10 muscles; *P* < 10^−9^, one-way ANOVA, \**P* < 0.0005, paired t-test). Error bars, SEM.

The present findings could offer an unprecedented perspective essential for understanding the functional role of the M1 in the volitional control of limb movements. There remains a long-standing controversy as to at what level the M1 is involved in movement control, i.e., low-level movement dynamics vs. high-level movement kinematics (*1, 2*). Considering that M1 activity affects muscle activity via two pathways with different time lags, we might obtain a comprehensive interpretation of why the wide range of tuning properties has been reported for M1 neurons. Neuronal discharges recorded in a short-time window are likely to represent movement dynamics because only the direct effect of the M1 on muscles is assessed. Conversely, neuronal discharges recorded in a long-time window are likely to represent movement kinematics due to assessment of the direct and trans-afferent effects on muscles. Future studies that reveal the causal relationship of the effect of the spinal reflex on muscle activity with the functional role of M1 neurons will help us understand how the central nervous system controls voluntary limb movements.

## Materials and Methods

### Animals

The dataset used in this study is the same dataset used in our previous study (*18*). We used one adult male monkey (monkey T, weight 6–7 kg) (*Macaca fuscata*) and one adult female monkey (monkey C, weight 5–6 kg) (*Macaca mulatta*). The animals were housed individually in temperature-controlled environments on a 12-h light/dark cycle. The experiments were approved by the experimental animal committee of the National Institute of Natural Sciences and the animals were cared for and treated humanely in accordance with National Institutes of Health guidelines.

### Behavioral task

The methods for the behavioral task, surgery, and recording of neuronal, muscular, and kinematic signals have been previously described (*18*). Briefly, both monkeys were operantly conditioned to perform a reach-to-grasp task with the right hand (Fig. 1A). After putting its hand on a home button for 2–2.5 s, each monkey reached for a lever and pulled it to receive a reward. We recorded the times for releasing the home button, pulling the lever, and pushing the home button. We analyzed data from 21 sessions of 10 min each trial per session for monkey T and 7 sessions for monkey C, in which the neuronal activity in peripheral afferents, electrocorticography (ECoG), electromyography (EMG), and kinematics were recorded simultaneously.

### Surgery

For EMG recording, we implanted pairs of Teflon insulated wire electrodes (AS631; Cooner Wire) into the forelimb muscles on the right side. We used the activity in the deltoideus posterior (Del), triceps brachii (Tri), biceps brachii (Bi), brachioradialis (BR), extensor carpi radialis (ECR), flexor carpi radialis (FCR), flexor carpi ulnaris (FCU), extensor digitorum communis (EDC), palmaris longus (PL), flexor digitorum superficialis (FDS), abductor pollicis longus (APL), and adductor pollicis (AP) of monkey T, and the Del, triceps brachii longus (TriLo), triceps brachii lateralis (TriLa), BR, ECR, FCU, EDC, flexor digitorum profundus (FDP), APL, and AP of monkey C.

We implanted a 32-channel grid electrode array (Unique Medical) with an electrode diameter of 1 mm and an interelectrode distance of 3 mm beneath the dura mater for ECoG of the sensorimotor cortex (Fig. 4A). We placed the ground and reference electrodes over the ECoG electrode so they were in contact with the dura.

We implanted two multielectrode arrays, consisting of 48 platinized-tip silicon electrodes with 0.1–1 MΩ at 1 kHz, a length of 1 mm, an interelectrode distance of 400 µm, and a 5 × 10 configuration (Blackrock Microsystems), into two cervical dorsal root ganglions (monkey T, C7 and C8; monkey C, C6 and C7) on the right side to record activity in peripheral afferents. We placed the reference wires over the dura of the spinal cord.

### Movement recordings

Forelimb movements were recorded with an optical motion capture system that used 12 infrared cameras (Eagle-4 Digital RealTime System; Motion Analysis). The spatial positions of the reflective markers (4- or 6-mm-diameter spheroids) were sampled at 200 Hz. Ten markers were attached to the surface of the forelimb as follows: left shoulder (marker 1, m1), center of the chest (m2), right shoulder (m3), biceps (m4), triceps (m5), lateral epicondyle (m6), medial to m6 (m7), radial styloid process (m8), ulnar styloid process (m9), and metacarpophalangeal joint of digit 2 (m10). We calculated flexion/extension (FE) of the shoulder, adduction/abduction (AA) of the shoulder, FE of the elbow, pronation/supination (PS), FE of the wrist, and radial/ulnar (RU) movement of the wrist (table S1). We applied a low-pass filter (5 Hz) to temporal changes in the joint angles to reduce noise.

### Neural recordings and spike detection

EMG signals were amplified using amplifiers (AB-611J; Nihon Kohden) with a gain of ×1,000–2,000 and sampled at 2,000 Hz in monkey T and 1,000 Hz in monkey C. We applied a 2nd-order Butterworth bandpass filter (1.5–60 Hz) to the signals. We rectified the filtered signals and performed resampling to 200 Hz. We smoothed the resampled signals using a moving window process with a window length of 11 bins.

ECoG signals were amplified using a multichannel amplifier (Plexon MAP system; Plexon) with a gain of ×1,000 and sampled from each electrode at 2,000 Hz in monkey T and 1,000 Hz in monkey C. We applied a 2nd-order Butterworth bandpass filter (1.5–240 Hz) to the signals. We computed a short-time fast Fourier transform on moving 100-ms windows of the filtered signals. We used a 200-Hz frequency step size. We computed power normalized to the average power in each session and calculated the average power in the high-gamma bands (high-gamma 1, 60–120 Hz; high-gamma 2, 120–180 Hz). We considered the high-gamma power of the ECoG signals as representative of neural activity in cortical areas in the analyses (*19*).

Neuronal signals of peripheral afferents were initially amplified using the same multichannel amplifier with a gain of ×20,000 and sampled from each electrode at 40 kHz. We extracted filtered waves (150–8,000 Hz) above an amplitude threshold that was determined by the “auto-threshold algorithm” of the software (Plexon MAP system). We sorted the thresholded waves using semiautomatic sorting methods (Offline Sorter; Plexon), followed by manual verification and correction of these clusters if needed. We removed a spike that occurred within 1 ms from a previous spike. We could isolate 25–39 units in monkey T, and 11–15 units in monkey C. To obtain the instantaneous firing rate, we convolved the inversion of the interspike interval with an exponential decay function whose time constant was 50 ms. We then resampled the firing rate to 200 Hz. By moving the forelimb, tapping over the muscle belly, and brushing the skin, we identified 2 units as muscle spindles, 1 unit as a tendon organ, 1 unit as a joint receptor, and 1 unit as a cutaneous receptor in monkey T and 1 unit as a muscle spindle, 1 unit as a tendon organ, and 3 units as cutaneous receptors in monkey C (*18*).

We calculated the movement-related modulation of the EMG signals, ECoG signals, and peripheral afferent activity before analyzing the data. We calculated the baseline activity by averaging the activity from -750 to -250 ms around movement onset. We then subtracted the baseline activity from the preprocessed activity. We used movement-related modulation throughout the premovement and movement periods (−500 to 1,000 ms around movement onset) as a single trial for further analysis.

### Sparse linear regression

We applied a Bayesian sparse linear regression algorithm that introduces sparse conditions for the unit/channel dimension only, and not for the temporal dimension of the model. Muscle activity was modeled as a weighted linear combination of high-gamma activity in the MCx and/or neuronal activity of peripheral afferents using multidimensional linear regression as follows:

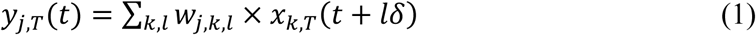

where, *y*_*j,T*_*(t)* is a vector of the EMG activity of muscle *j* (12 and 10 muscles in monkey T and C, respectively) at time index *t* in trial *T*; *x*_*k,T*_*(t+lδ)* is an input vector of peripheral afferent or cortical signal *k* at time index *t* and lag time *lδ* (*δ* = 5 ms) in trial *T*; and *w*_*j,k,l*_ is a vector of weights on the peripheral afferent or cortical signal *k* at lag time *lδ*. As we examined how the combined activity in the MCx and/or peripheral afferents influenced muscle activity, lag time *lδ* (Equation 1) was set to negative values. For the direct effect of the MCx on muscle activity via the descending pathway (descending input), we used activity in the MCx from -50 to -5 ms to reconstruct muscle activity at time 0 for the following reasons. Averaging the muscle activity triggered at the spiking activity of single neurons in the M1 shows postspike facilitation with a latency of 6.7 ms (*8*). Accordingly, we set 5 ms as the shortest lag time. The weighted sum of M1 neuronal activity accurately accounted for muscle activity at a lag time of 40–60 ms (*20*). Conversely, there is a possibility that the MCx has some effects on muscle activity via the spinal reflex arc within 60–70 ms, as shown in fig. S4. To avoid the influence of the spinal reflex arc, we set 50 ms as the longest lag time. For the effect of peripheral afferents on muscle activity (afferent input), we used the activity in the peripheral afferents from -50 to -5 ms to reconstruct muscle activity at time 0 for the following reasons. Averaging the muscle activity triggered at the spiking activity of peripheral afferents shows postspike facilitation with a latency of 5.8 ms (*21*). Thus, we set 5 ms as the shortest lag time. To avoid the influence of the spinal reflex arc, as described above, we set 50 ms as the longest lag time. For the reafferent effect of the MCx on muscle activity via the spinal reflex arc (trans-afferent input), we used the activity in the MCx from -65 to -110 ms for monkey T and from -75 to -120 ms for monkey C to reconstruct muscle activity at time 0. We obtained this lag time by simply adding the time when the MCx most accurately accounted for the activity of the peripheral afferents (60 ms for monkey T and 70 ms for monkey C) to the lag time of the descending input. To examine the contribution of the descending and afferent inputs on muscle activity within short time windows, we built a model to reconstruct muscle activity from a weighted linear combination of descending and afferent inputs within an overlapping, sliding time window of 500 ms.

To compute the contribution of each descending, afferent, and trans-afferent input to the reconstruction of muscle activity, we calculated each component of reconstructed activity using either input and the respective weight values in a decoding model that was built from the combined inputs. For example, the descending component was calculated as follows:

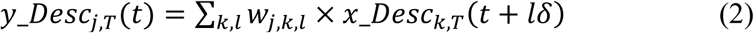

where, *y_Desc*_*j,T*_(*t*) is a vector of the descending component at muscle *j* at time index *t* in trial *T*; *x_Desc*_*k,T*_(*t+lδ*) is an input vector of cortical signal *k* at time index *t* and lag time *lδ* in trial *T*; and *w*_*j,k,l*_ is derived from a vector of weights in Equation (1), but with weights assigned to peripheral afferents removed.

The activity of peripheral afferents was modeled as a weighted linear combination of neuronal activity in the high-gamma power band in the MCx using multidimensional linear regression, as follows:

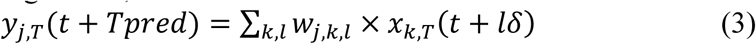

where, *y*_*j,T*_*(t)* is a vector of peripheral afferent *j* at time index *t* in trial *T*; *x*_*k,T*_*(t+lδ)* is an input vector of cortical signal *k* at time index *t* and lag time *lδ* (*δ* = 5 ms, *l* = -10–-1) in trial *T*; and *w*_*j,k,l*_ is a vector of weights on cortical signal *k* at lag time *lδ*. We considered that neuronal activity in the MCx evokes muscle activity, which in turn generates the activity of peripheral afferents. We set lag time *lδ* (Equation 3) to negative values. By changing *Tpred* from 0 ms to 120 ms, we attained *Tpred*, at which the accuracy of reconstructing the activity in peripheral afferents was the largest (60 ms for monkey T and 70 ms for monkey C, fig. S4D).

### Data analysis

We built models to reconstruct the temporal changes in the EMG signals or activity in peripheral afferents using a partial dataset (training dataset) and tested them using the remainder of the same dataset (test dataset). One hundred and eight trials were selected randomly as a training dataset, and 21 trials were selected randomly from the remainder as the test dataset. To assess the model, we calculated the correlation coefficient between the observed data and their reconstruction in the test dataset. We also calculated the variance accounted for (VAF) as follows:

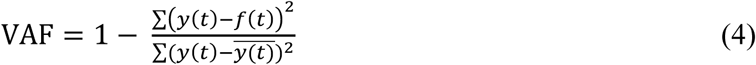

where, *y*(*t*) is a vector of the actual activity in muscles at time index 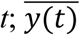 is the mean of *y(t)*; and *f(t)* is the reconstructed activity at time index *t*. We performed 6-fold cross-validation in the analysis of each session and used averaged values for the analysis. Then, we calculated the averaged values of each muscle or peripheral afferent from data taken from 21 (monkey T) and 7 (monkey C) sessions. In control analyses of the model reconstruction, we created surrogate training datasets in which we shuffled the temporal profiles of the inputs independently across different blocks to generate a model, and subsequently tested the model.

To obtain the onset time of the observed muscle activity or the reconstruction, we first calculated the average of the aligned waveform in the test dataset. Then, we defined one-fifth of the maximum amplitude of the observed muscle activity from 250 ms before to 250 ms after movement onset as a threshold. If the activity or the reconstruction was over the threshold in 5 consecutive bins, the first of these bins was set as the onset. The calculated onset corresponded well with that based on visual inspection. We calculated the average onset values from six test datasets in one session and finally obtained their average from all sessions. We calculated the average positive or negative values of each component during a period in which movement-related modulation of muscles was observed (−100 to 1,000 ms around movement onset). We then normalized the averaged values of each component by the average positive or negative values of the reconstruction from the combined inputs during this period (Fig. 3, and fig. S6). We tested whether the normalized positive or negative values deviated from zero using a standard t-test. The time of the initial peak of the wrist joint angle along the FE direction of monkey T was 51.4 ± 9.6 ms (mean ± standard deviation (SD)) and those of the wrist joint angle along the FE direction and elbow joint angle along the PS direction of monkey C were 31.4 ± 5.6 ms and 47.1 ± 2.7 ms, respectively. We calculated the mean time value of each component during a period from the beginning of the reaching movement (from 50 to 100 ms around movement onset) (fig. S3). We tested whether the mean time values deviated from zero using a standard t-test.

We examined weight values assigned to four types of cortical input: descending input in a model that reconstructed muscle activity from descending and afferent inputs; motor cortical input in a model that reconstructed afferent activity from activity in the MCx; descending input in a model that reconstructed muscle activity from descending and trans-afferent inputs; and trans-afferent input in a model that reconstructed muscle activity from descending and trans-afferent inputs (Fig. 4A). To obtain a weight value for each electrode, the absolute weight vector *w*_*j,k,l*_ in Equation (1) or (3) was averaged across time points. Values for high-gamma 1 and 2 were averaged. One*-*way ANOVA was used to determine whether there were any statistically significant differences between the means of the weight values of different electrodes.

### Statistical analysis

We used the nondirectional paired or unpaired Student’s *t*-test. When comparing more than 2 group means, we first assessed the data using one*-*way ANOVA. An alpha level of significance was set at 0.05 for all statistical tests. Data are expressed as the mean ± standard error of the mean (SEM) or the mean ± SD. We used MATLAB R2015b (MathWorks) for the statistical analysis. The data distribution was assumed to be normal, but this was not tested formally. No statistical methods were used to predetermine sample sizes. However, sample sizes were estimated by methodologically comparable previous experiments in our laboratory and are similar to those employed in the field.

## Acknowledgments

We thank Y. Yamanishi for animal care, Y. Nishihara, M. Suzuki, and M. Togawa for technical help, and Professor K. Seki for encouragement.

## Author contributions

T.U. and Y.N. designed and performed the experiments. T.U. analyzed the data and prepared the figures. T.U., T.I., and Y.N. wrote the manuscript.

## Data and materials availability

All data needed to evaluate the conclusions in the paper are present in the paper and/or the Supplementary Materials. Additional data related to this paper may be requested from the authors.

