## Supplemental file for "The primary motor cortex prospectively computes the spinal reflex"

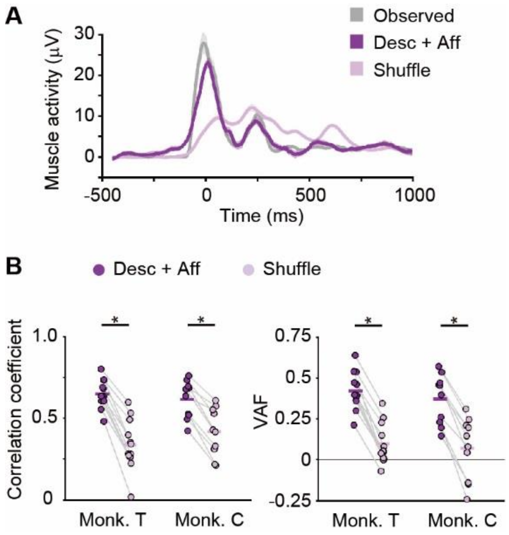


**Fig. S1. Descending and afferent inputs account for muscle activity.** (**A**) Average modulation of the observed muscle activity, reconstruction using descending and afferent inputs, and shuffled control data aligned to movement onset. Shaded areas, SEM. (**B**) Mean reconstruction accuracy. Correlation coefficient and VAF between the observed and reconstructed traces (monkey T, n = 12 muscles; monkey C, n = 10 muscles; *$P < 0.0001$, paired t-test). Superimposed bars, mean.

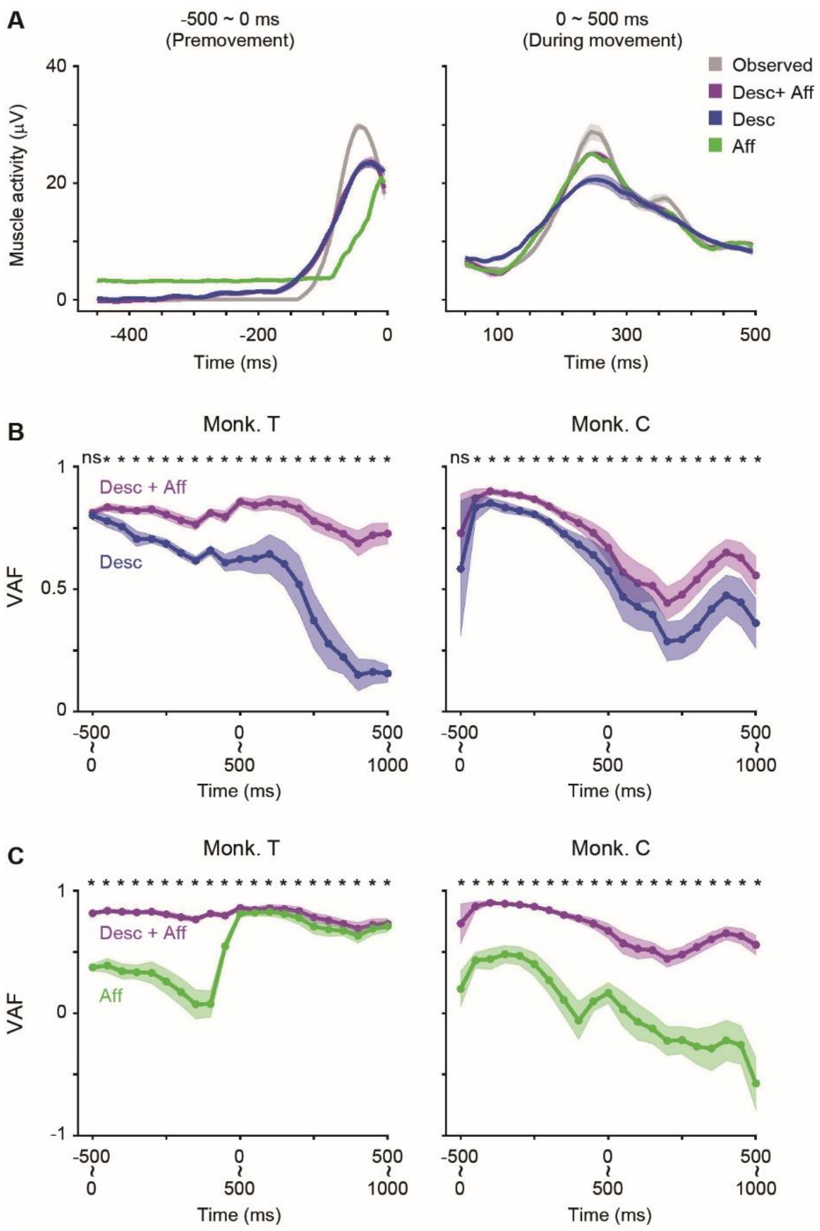


**Fig. S2. Descending, not afferent, input contributes to the reconstruction of muscle activity before movement onset.** (**A**) Average modulation of the observed muscle activity, reconstruction using descending and afferent inputs, and each activity in two 500-ms time windows (premovement period, -500 to 0 ms around movement onset; during movement, 0 to 500 ms around movement onset). (**B, C**) Mean reconstruction accuracy between the observed and reconstructed traces for each 500-ms sliding window (B, Desc + Aff vs. Desc; C, Desc + Aff vs. Aff) (monkey T, n = 12 muscles; monkey C, n = 10 muscles; *$P < 0.05$, paired t-test). ns, not significant. Shaded areas, SEM. In the premovement period, no significant difference was detected in reconstruction accuracy between models built from descending and afferent inputs and models built from descending input only.

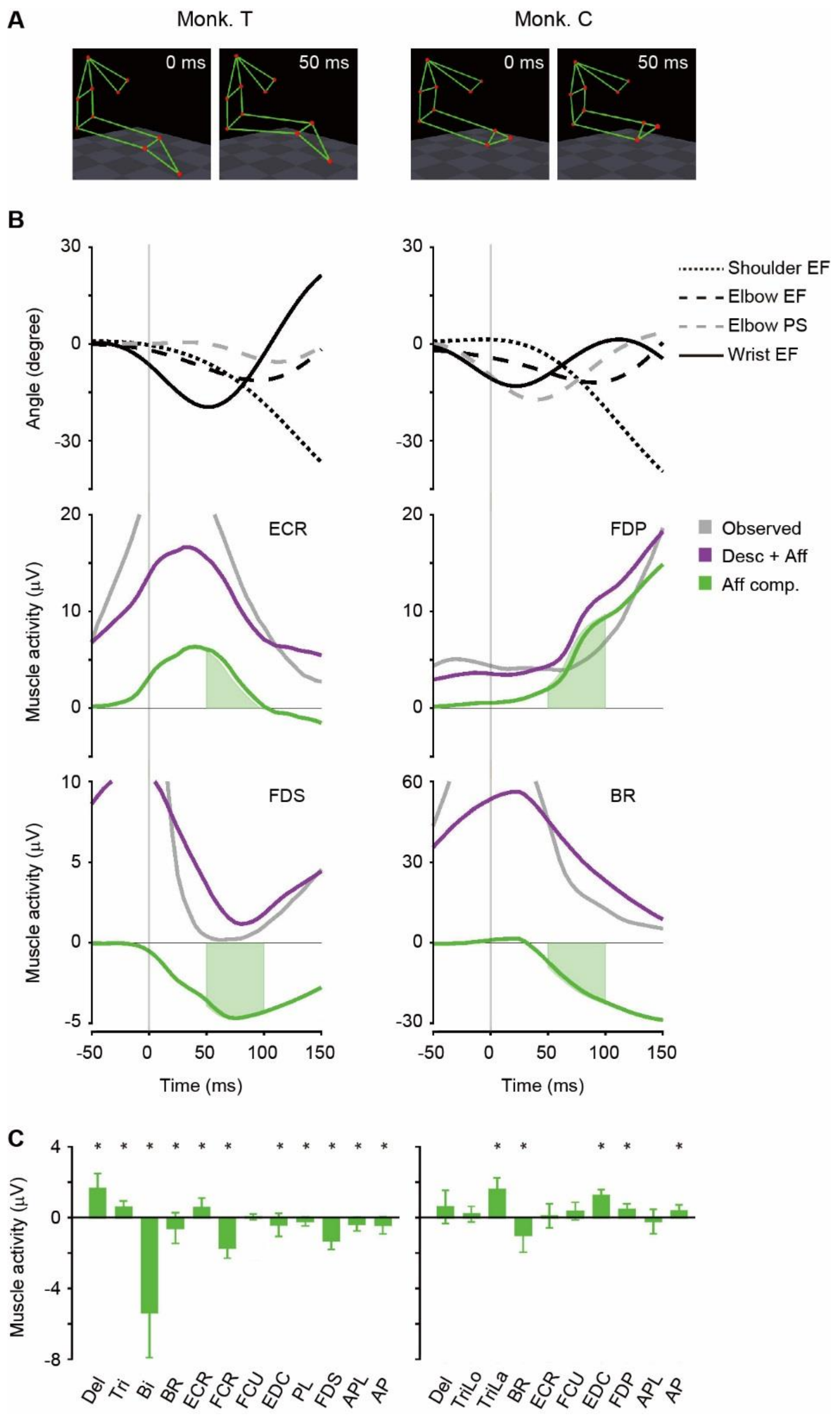


**Fig. S3. The effects of afferent input on muscle activity are partly accounted for by spinal reflex action.** (**A**) Stick figures of the forelimb and chest when the monkeys began to reach. Monkey T exhibited wrist flexion and monkey C performed elbow supination and wrist flexion. (**B**) *Top*: Four forelimb joint angles. *Second* and *Third*: Average modulation of the observed muscle activity, reconstruction using descending and afferent inputs, and afferent component in the reconstruction. Vertical lines, time of movement onset. (**C**) The mean time value of modulation of afferent components for each muscle in a period from the beginning of the reaching movement (50 to 100 ms around movement onset; shown in the green area in fig. S3B) (monkey T, n = 21 sessions; monkey C, n = 7 sessions; *$P < 0.05$, t-test). Error bars, SD.

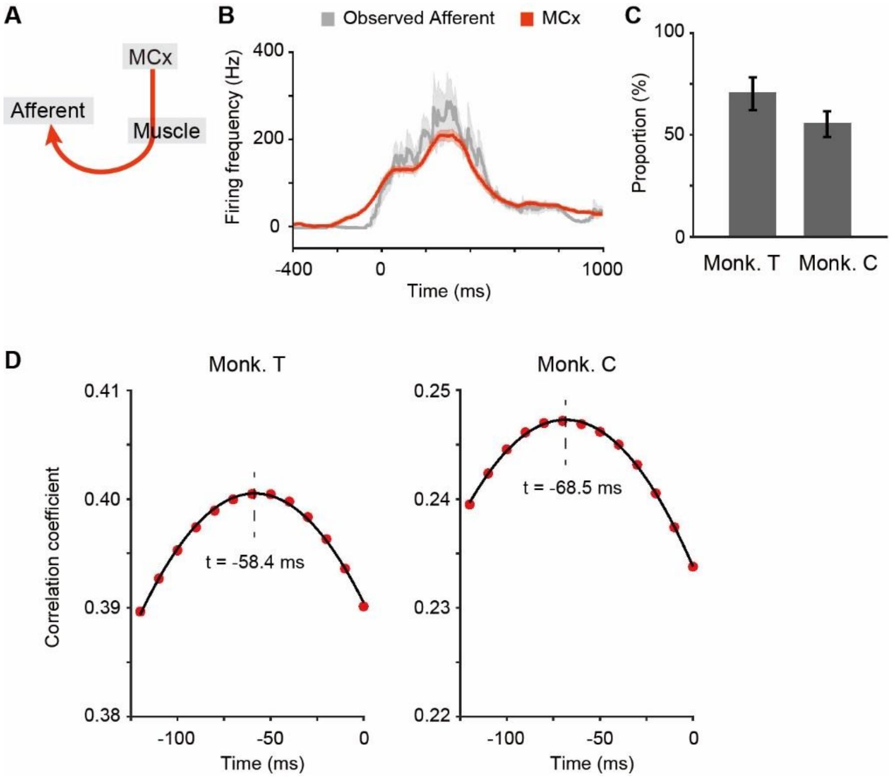


**Fig. S4. MCx activity accounts for subsequent afferent activity.** (**A**) Model to account for afferent activity from MCx activity. (**B**) Average modulation of the observed afferent activity and reconstruction using activity in the MCx aligned to movement onset. Shaded areas, SEM. (**C**) Proportion of afferents whose activity was reconstructed accurately from MCx activity more than models built from the shuffled data (monkey T, n = 21 sessions; monkey C, n = 7 sessions). MCx activity could explain the activity of more than 50% of peripheral afferents in a linear model. Bar graphs and error bars, mean ± SEM. (**D**) Correlation coefficients between the observed and reconstructed traces were plotted against the end of lag times of a 50-ms window between afferent and MCx activity. The plot was fitted by the quadratic curve shown with a black line. Vertical dotted lines represent the lag time at which the maximum value was obtained on the fitted curve.

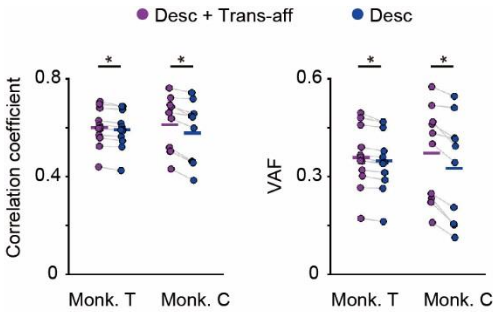


**Fig. S5. Trans-afferent input is important for the reconstruction of muscle activity.** Mean reconstruction accuracy of the model obtained from descending and trans-afferent inputs and descending input only. Correlation coefficient and VAF between the observed and reconstructed traces (monkey T, n = 12 muscles; monkey C, n = 10 muscles; *$P < 0.0005$, paired t-test). Superimposed bars, mean.

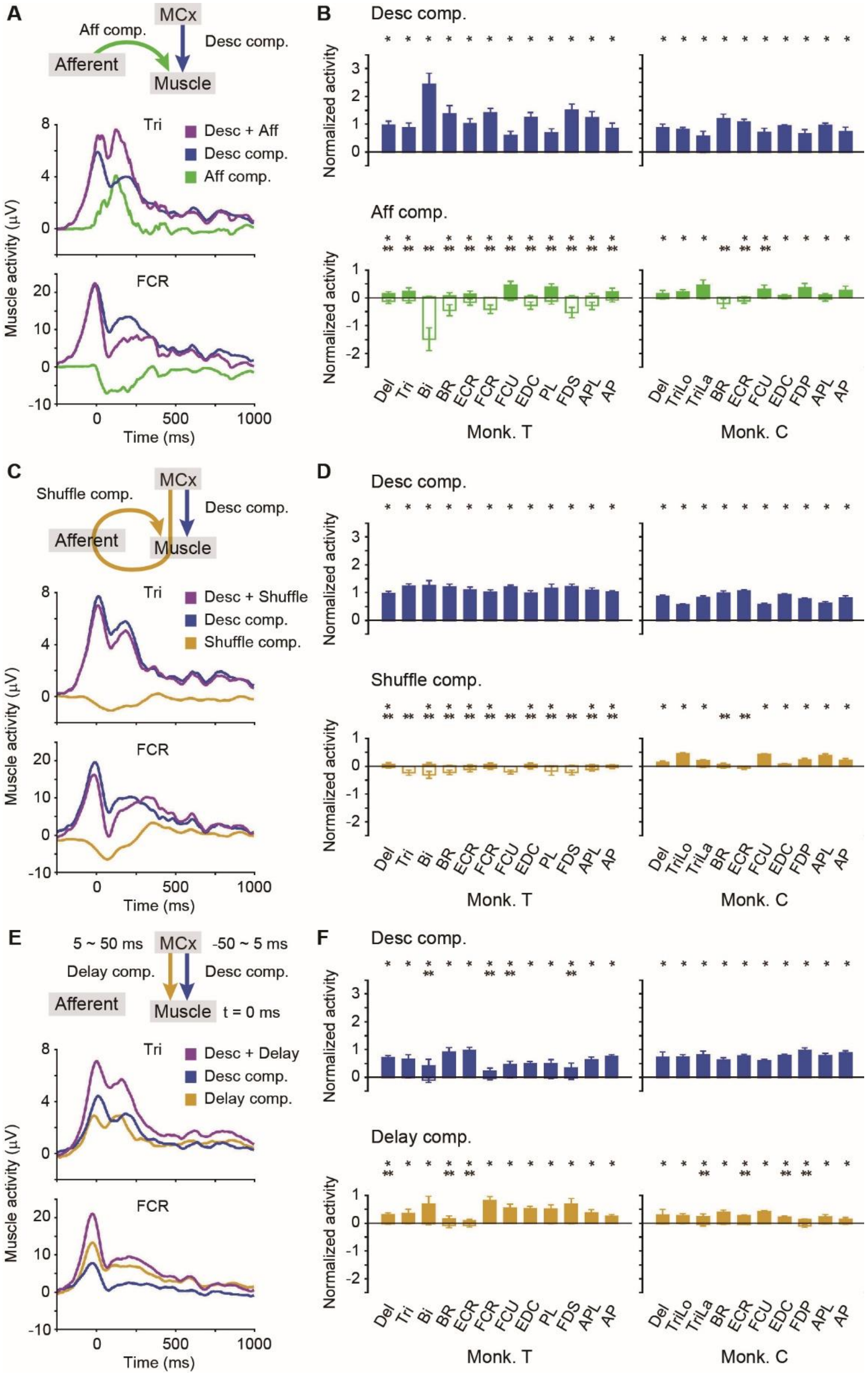


**Fig. S6. Control models fail to duplicate the corresponding component obtained by a model built from descending and afferent inputs.** (**A**) Average modulation of the reconstruction of a shoulder muscle (triceps) and wrist muscle (FCR) using descending and afferent inputs, and each component in the reconstruction aligned to movement onset. (**B**) Modulation of descending and afferent components for each muscle normalized by the total modulation of the reconstruction from the combined inputs from -100 to 1,000 ms around movement onset (monkey T, n = 21 sessions; monkey C, n = 7 sessions; *$P < 0.05$, unpaired t-test for positive values, **$P < 0.05$, unpaired t-test for negative values). Error bars, SD. (**C, D**) Same as (A, B) for descending and shuffled data from trans-afferent inputs. (**E, F**) Same as (A, B) for descending input and delayed activity in the MCx.

| Joint angle | Two vectors | |
|---|---|---|
| shoulder FE | cross product of a vector from m3 to m2 and one from m3 to m1 | vector from m3 to m7 |
| shoulder AA | vector from m3 to m1 | vector from m3 to m7 |
| elbow FE | vector from m7 to m3 | vector from m7 to the center of m8 and m9 |
| elbow PS | projection of a vector from m7 to m3 on a plane with a normal vector from m7 to the center of m8 and m9 | projection of a vector from m8 to m9 on the same plane |
| wrist FE | cross product of a vector from m8 to m10 and one from m8 to m9 | vector from the center of m8 and m9 to m7 |
| wrist RU | vector from m9 to m8 | vector from the center of m8 and m9 to m10 |

**Table S1. Calculation of joint angles.** Joint angles were calculated from the two vectors presented in the right two columns. In particular, Euler angles are used to represent relative joint rotations.
